# *Aspergillus latus*: a cryptic causative agent of aspergillosis emerging in Japan

**DOI:** 10.1101/2024.11.25.625089

**Authors:** Saho Shibata, Momotaka Uchida, Sayaka Ban, Katsuhiko Kamei, Akira Watanabe, Takashi Yaguchi, Vit Hubka, Hiroki Takahashi

## Abstract

Allodiploid hybrid species, *Aspergillus latus*, belonging to section *Nidulantes*, is a hybrid of *A. spinulosporus* and unknown species closely related to *A. quadrilineatus* and *A. subltus*. This hybrid has often been misidentified as the species in section *Nidulantes*, such as *A. nidulans*, *A. spinulosporus*, *A. sublatus*, or other cryptic species*. A. latus* has not been reported in Japan as well as Asia so far. In this study, we screened 23 clinical strains identified as *A. spinulosporus* isolated in Japan from 2012 to 2023 and found seven *A. latus* strains. To characterize the *A. latus* strains, we conducted comprehensive phenotyping including morphological observation, whole genome sequences and phylogenetic analysis based on calmodulin (*CaM*) gene. In addition, we conducted antifungal susceptibility testing for *A. latus* strains. As a result, the morphological characters of *A. latus* were more similar to those of *A. spinulosporus* compared to *A. sublatus*. However, the ascospore of *A. latus* differed from that of *A. spinulosporus*. *A. latus* phylogenetically clustered with *A. spinulosporus* and *A. sublatus*. Furthermore, *A. latus* strains showed reduced susceptibility to caspofungin and amphotericin B compared to *A. spinulosporus*, while they were susceptible to azoles. Our results suggest that *A. latus* has been a causative pathogen of aspergillosis in Japan since 2013.

## Introduction

*Aspergillus* species are the causal agents for aspergillosis, mainly affecting immunocompromised individuals [1, 2]. Although *A. fumigatus* is a main causative agent, other species such as *A. nidulans*, *A. flavus*, *A. niger*, and *A. terreus* also contribute to the disease [3–6]. *Aspergillus nidulans*, a member of the section *Nidulantes* poses a serious threat to immunocompromised patients, especially those with chronic granulomatous disease [3]. Additionally, eleven other species within the section *Nidulantes* have been isolated from patients with invasive aspergillosis [7]. Recently, Steenwyk et al. (2020) through screening of *A. nidulans* strains identified that *A. latus* is an allodiploid hybrid species between *A. spinulosporus* and unknown species closely related to *A. quadrilineatus*, and exhibits higher virulence and minimum inhibitory concentrations to some antifungals than *A. nidulans* [8]. This highlights the importance of accurate species identification to ensure effective treatment, as misidentification could compromise patient care.

Steenwyk et al. (2020 & 2024) reported 30 *A. latus* strains from Belgium, Brazil, France, Germany, Netherlands and Portugal identified as *A. nidulans*, *A. spinulosporus*, or other cryptic species, suggesting that *A. latus* infections are likely underreported [9]. Prospective survey in single center in Japan reported that six *A. nidulans* strains were identified among 107 *Aspergillus* strains [11]. To date, *A. latus* has not been reported in Asia including Japan. However, *A. nidulans* and some cryptic species were isolated from patients in Japan, namely *A. latus* could had also been isolated [10, 11].

In this study, we identified seven clinical *A. latus* strains isolated in Japan by screening 23 *A. spinulosporus* strains which were collected between 2012 and 2023. To characterize these seven *A. latus* strains isolated in Japan, we performed comprehensive phenotyping including morphological observations, whole genome sequences and phylogenetic analysis based on calmodulin (*CaM*) gene. In addition, we measured the antifungal susceptibility of these strains to six antifungals. Here, we present the first report of *A. latus* being isolated in Japan. Our findings indicate that *A. latus* has been the causative agent of aspergillosis in Japan since 2013.

## Material and methods

### Strains

*A. latus*, *A. sublatus* and *A. spinulosporus* strains used in this study have been stored and maintained at the Medical Mycology Research Center, Chiba University (IFM strains) in Japan. A total of 23 strains of *A. latus* and *A. spinulosporus* are clinical isolate in Japan, and *A. sublatus* IFM 42029 (= IFM 4553 = CBS 140630) is a type strain collected from soil in Brazil.

### Genomic DNA extraction

Fungi were incubated on Potato Dextrose Agar (Thermo Fischer Scientific, MA, USA) at 30 °C for 7 days to obtain fully mature conidia. The conidia were inoculated in Potato Dextrose Broth (Thermo Fischer Scientific) at 37 °C at 180 rpm for 16 h. Mycelia were washed with H_2_O, and genomic DNA was purified with phenol-chloroform extraction and Monarch^®^ Genomic DNA Purification Kit (New England Biolabs, MA, USA) [12, 13]. The DNA concentration and quality were then measured by Qubit Fluorometer (Thermo Fischer Scientific). The extracted DNA were used for cloning of *CaM* gene and whole-genome sequencing.

### Whole genome sequencing and assembly

Whole-genome 150 bp paired-end (PE) sequencing of four *A. latus* strains, IFM 65030, IFM 65233, IFM 65239 and IFM 66778 was performed using Illumina NovaSeq 6000 (Illumina, CA, USA) by Novogene Biotech Co. (Beijing, China). The raw genomic reads were trimmed using fastp v.0.20.1 [14]. The mitochondrial genomes were assembled using GetOrganelle v.1.6.4 [15]. To filter the mitochondrial reads, the reads were aligned with the mitochondrial genome using BWA-MEM v.0.7.17-r1188 [16]. The mapped reads were filtered using SAMtools v.1.10 [17] and SeqKit v.0.10.1 [18]. The nuclear genomes were assembled using VelvetOptimiser v.2.2.6 [19].

The completeness of draft genomes was evaluated by BUSCO v. 5.2.1 [20] with the database eurotiales_odb10. blastn v.2.15.0+ [21] was used for extracting the sequences of *CaM* of *A. latus* strains by serving the sequence of *A. nidulans* FGSC A4 as query sequence. The genome sizes of *A. latus* strains were estimated using Jellyfish [22] and GenomeScope with 21 k-mers [23].

### DNA Cloning and Sequencing

The cloning of two *CaM* genes of three strains, IFM 61956, IFM 63852 and IFM 64630, were performed. PCR amplification was performed by Quick Taq® HS DyeMix (TOYOBO, Tokyo, Japan) in a ProFlex™ PCR System (Thermo Fisher Scientific) with a program consisting of an initial denaturing step at 94 °C for 2 min: 35 cycles of denaturation at 94 °C for 30 sec, annealing at 50 °C for 30 sec, and extension at 68 °C for 1 min. The amplicons of *CaM* were obtained by PCR with primer pairs CF1L_SacI (5′-AAAGAGCTCGCCGACTCTTTGACYGARGAR-3′) and CF4_SacI (5′-AAAGAGCTCTTTYTGCATCATRAGYTGGAC-3′). These primers contain the sequence of CF1 and CF4 primers, respectively, from a previous study [24, 25], with the SacI restriction site at the 5′ end. The PCR amplicons were purified by gel extraction with QIAquick Gel Extraction Kit (QIAGEN, Hilden, Germany). The objective gene fragments were cloned into pMK-dGFP vector [26], and introduced into competent cells of *Escherichia coli* DH5α. FastGene Plasmid Mini Kit (Nippon Genetics, Tokyo, Japan) was used to extract the plasmid DNA. The plasmid (ca. 2.0 µL) was used as the templates for the sequencing reaction with a BigDye Terminator v3.1 Cycle Sequencing Kit (Thermo Fisher Scientific) along with PtrpC-Rev (5′-AAATGCTCCTTCAATATCATCTTCTGTCGA-3′) and M13R (5′-CAGGAAACAGCTATGAC-3′) primers [27], following the manufacturer’s instructions. Ethanol precipitation was used to remove excess fluorescent dye-terminators from cycle sequencing reactions prior to analysis on an automated sequencer. Sequence was performed with an ABI 3100 Genetic Analyzer (Thermo Fisher Scientific).

### Phylogenetic analysis

The sequences of *CaM* ranging from 438 bp to 824 bp in length were aligned with MAFFT v.7.508 [28, 29]. Aligned sequences were trimmed using trimAl v.1.4.rev15 [30], resulting in 686 bp in length. The maximum likelihood tree of *CaM* was constructed using the multithreaded version of RAxML v.8.2.12 [31], the GTRCAT model, and 1,000 bootstrap replicates. The phylogenetic tree was visualized using the FigTree (free download available at http://tree.bio.ed.ac.uk/software/figtree/). A total of 112 taxa were included in our tree (Supplementary Table S1).

### Morphological observations

*A. latus* IFM 65329 was used for microscopic and macroscopic observation. The colony character of the other six strains of *A. latus*, *A. sublatus* IFM 42029^T^ (= IFM 4553^T^) and *A. spinulosporus* IFM 66771 were observed. The observations of macroscopic characters were curried on the agar media, i.e., Czapek yeast autolysate agar (CYA, Czapeck Dox Agar, Duchefa Biochemie, Haarlem, Netherlands and Difco™ Yeast Extract, Thermo Fischer Scientific), oatmeal agar (OA, Sigma Aldrich, MO, USA) and malt extract agar (MEA, Thermo Fischer Scientific) with trace elements (0.1 g ZnSO_4_·7H_2_O and 0.5 g CuSO_4_· 5H_2_O in 100 mL distilled water) at 25 or 37 °C for 7 days. Microscopic observations of conidia and conidiophores were made from 1 week old colonies on MEA. Ascomata and ascospores were observed from 8-week-old colonies on OA. A light microscope (Nikon ECLIPSE Ni, Tokyo, Japan and Swift Optical Instruments S7-TP520, TX, USA) was used to examine morphological characters including the size and shape of ascomata, ascospores, conidiophores and conidia. These were mounted in a drop of water on glass slides. The slide preparations were examined and photographed using a digital camera (Nikon DS-Fi3, Japan). Approximately 30 structures were randomly chosen, and their length and width were measured using ‘ImageJ’ software (free download available at http://rsbweb.nih.gov/ij/).

### Antifungal susceptibility testing

Minimum inhibitory concentration (MIC) tests for amphotericin B (AMPH-B), itraconazole (ITCZ), voriconazole (VRCZ), miconazole (MCZ), micafungin (MCFG), and caspofungin (CPFG), were performed using the dried plate for antifungal susceptibility testing of yeasts method (Eiken Chemicals, Tokyo, Japan). The method follows the Clinical and Laboratory Standards Institute M38-E2 with slight modifications [32–34].

## Results

### Screening and identification of *A. latus*

To identify the presence of allodiploid hybrid strains of *A. latus*, we sequenced the *CaM* genes of 23 *A. spinulosporus* strains. Sequences with double peaks were observed in seven strains. The sequences of two copies of *CaM* were determined in all seven strains by either cloning or whole-genome sequencing (Table 1). These strains were isolated from the patients between 2013 and 2022. The ages of the patients ranged from 13 to 79 years. The patients were diagnosed with invasive pulmonary aspergillosis (IFM 63852 and IFM 66778) and chronic pulmonary aspergillosis (IFM 65233 and IFM 65329), the other patients were not diagnosed with aspergillosis (IFM 61956, IFM 64360, and IFM 65030). However, the patients were diagnosed with other diseases such as leukemia. Therefore, we could not conclude that *A. latus* is a primary cause of death.

**Table 1.**
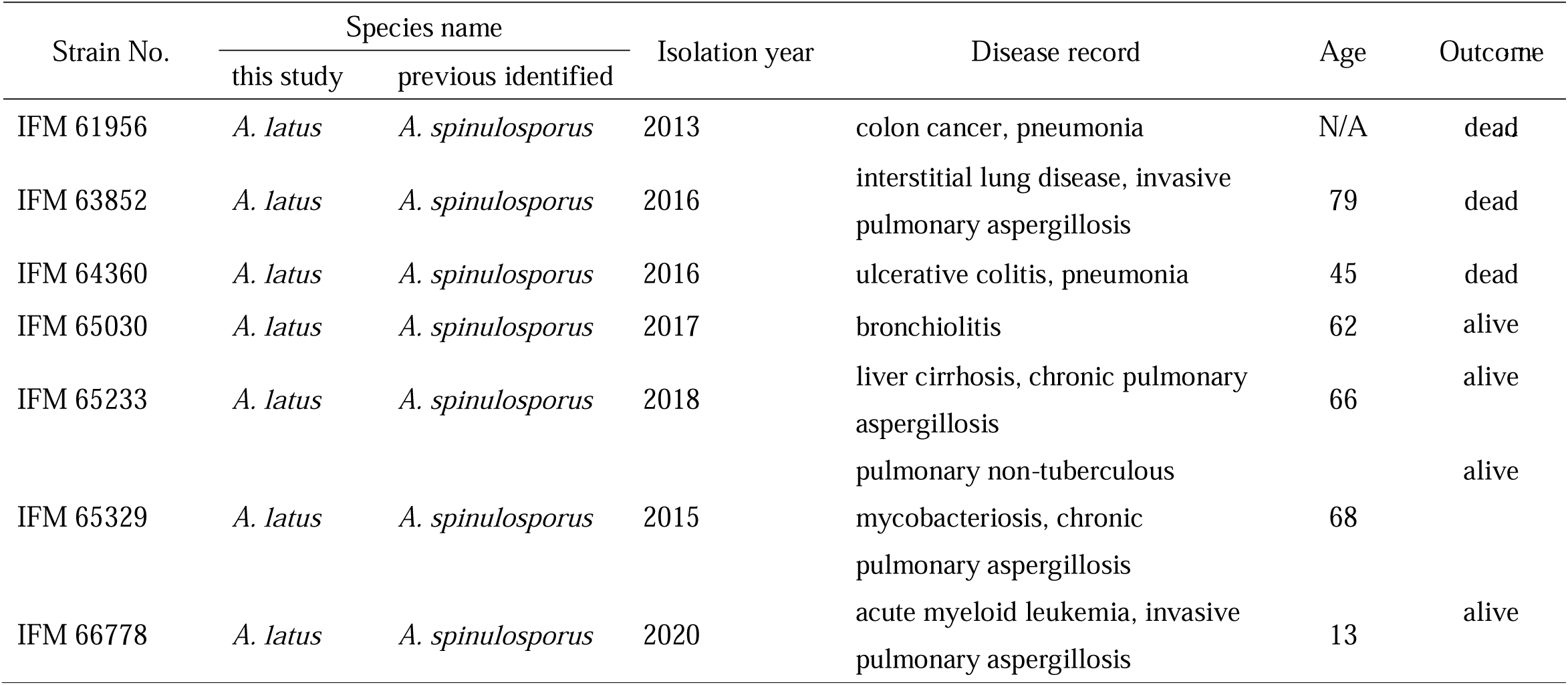
Clinical information of *Aspergillus latus* strains examined in this study.

| Strain No. | Species name |  | Isolation year | Disease record | Age | Outcome |
| --- | --- | --- | --- | --- | --- | --- |
|  | this study | previous identified |  |  |  |  |
| IFM 61956 | *A. latus* | *A. spinulosporus* | 2013 | colon cancer, pneumonia | N/A | dead |
| IFM 63852 | *A. latus* | *A. spinulosporus* | 2016 | interstitial lung disease, invasive pulmonary aspergillosis | 79 | dead |
| IFM 64360 | *A. latus* | *A. spinulosporus* | 2016 | ulcerative colitis, pneumonia | 45 | dead |
| IFM 65030 | *A. latus* | *A. spinulosporus* | 2017 | bronchiolitis | 62 | alive |
| IFM 65233 | *A. latus* | *A. spinulosporus* | 2018 | liver cirrhosis, chronic pulmonary aspergillosis | 66 | alive |
| IFM 65329 | *A. latus* | *A. spinulosporus* | 2015 | pulmonary non-tuberculous mycobacteriosis, chronic pulmonary aspergillosis | 68 | alive |
| IFM 66778 | *A. latus* | *A. spinulosporus* | 2020 | acute myeloid leukemia, invasive pulmonary aspergillosis | 13 | alive |

Draft genomes for four *A. latus* strains (IFM 65030, IFM 65233, IFM 65239 and IFM 66778) were generated, with assembled genome sizes ranging from 62.1 Mb to 63.2 Mb, aligning with the typical genome sizes of *A. latus* (previous study showed average genome size 69.09 ± 5.68 Mb [8]) (Supplementary Table S2).

### Phylogenetic analysis

We conducted a phylogenetic analysis to characterize the position of seven Japanese *A. latus* strains. One copy of *CaM* gene sequence clustered with the sequences of *A. spinulosporus*, while the other copy clustered with *A. sublatus* (Fig. 1). The *CaM* sequence of *A. latus* differed from that of *A. sublatus* in length by 1 bp, that of *A. quadrilineatus* in length by 6 bp, but the *CaM* sequence of *A. latus* harbors no specific substitution compared to *A. sublatus*, *A. quadrilineatus* and *A. spinulosporus*.

**Fig. 1.**
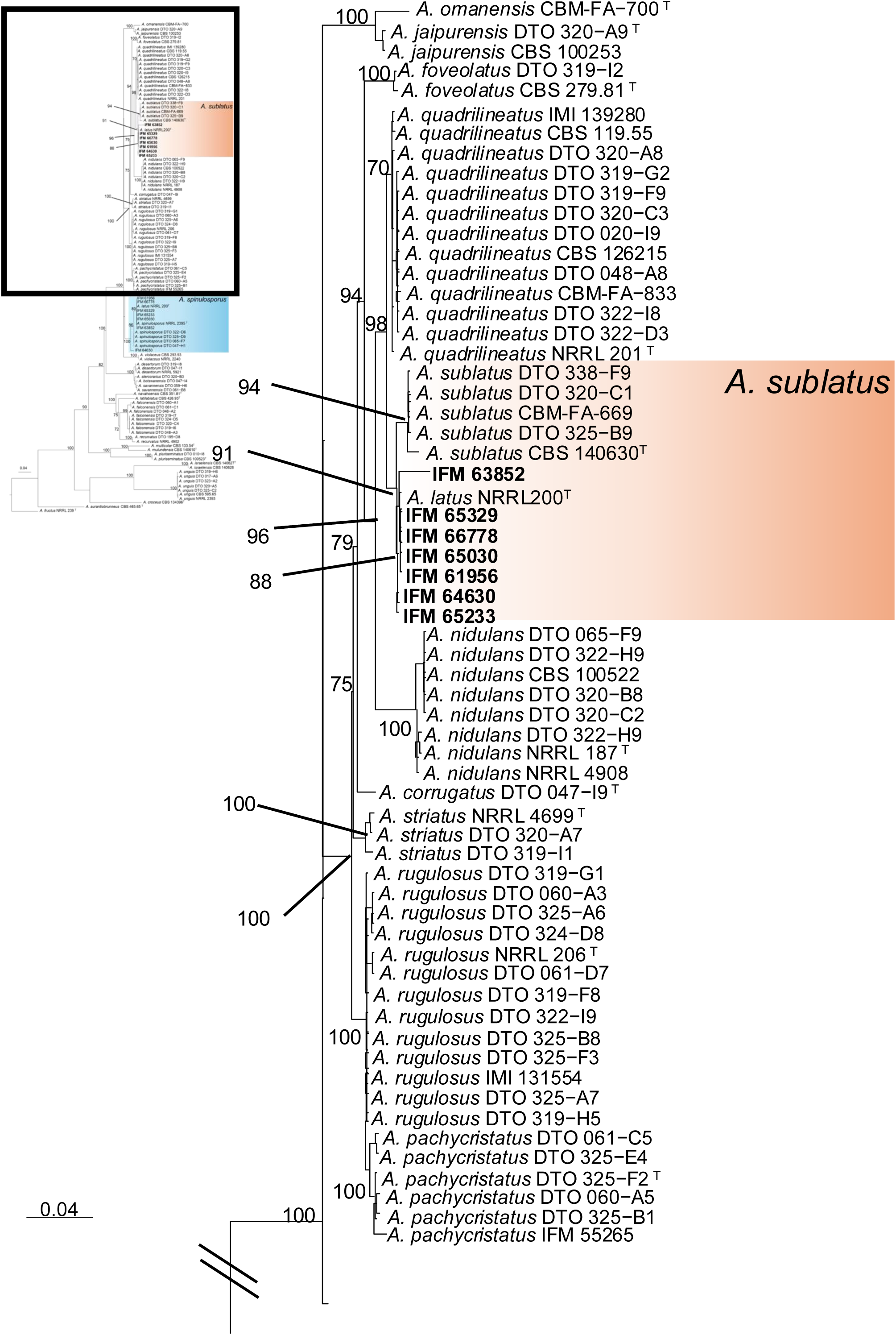
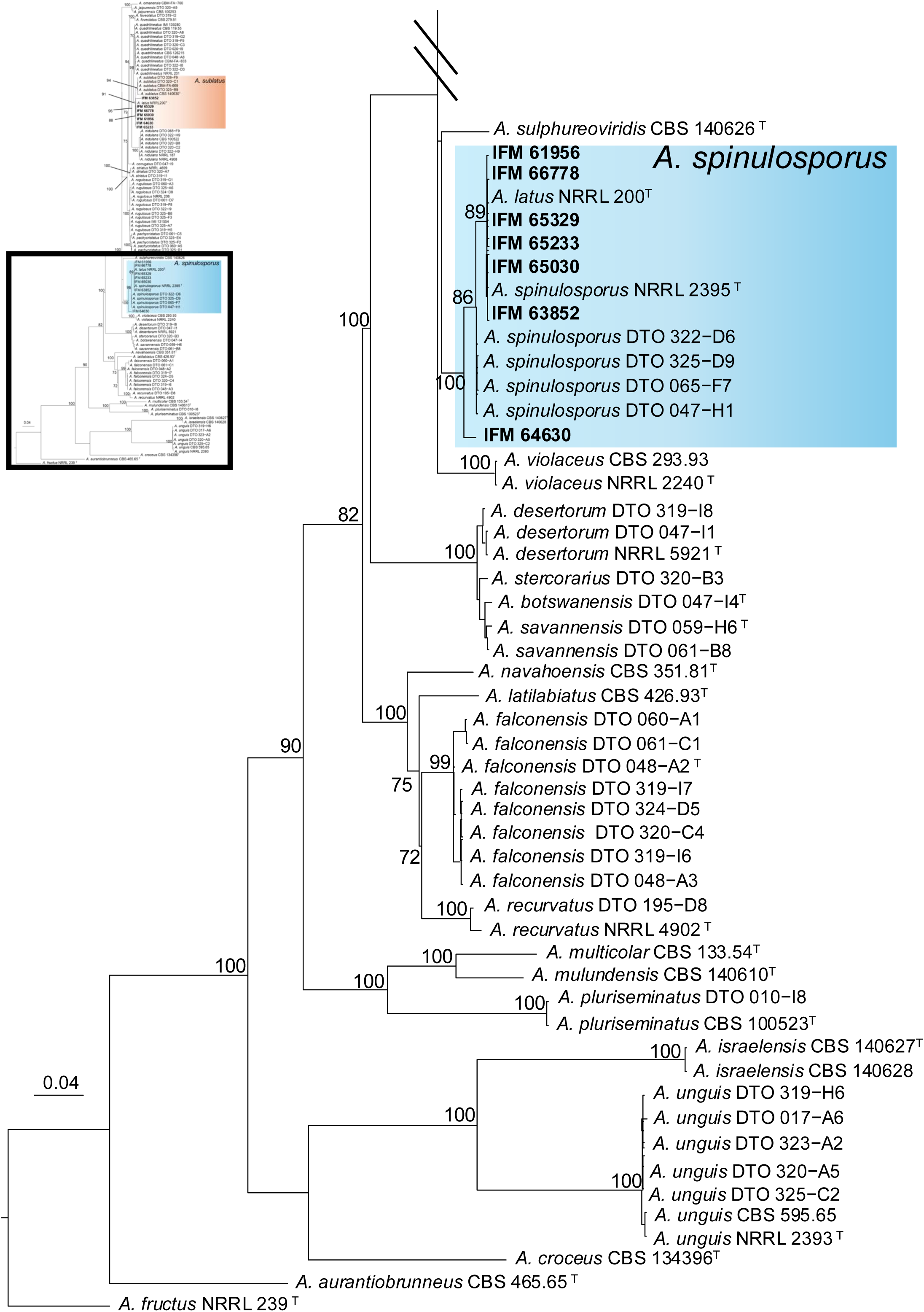
Phylogenetic tree constructed using maximum likelihood (ML) analysis based on the models selected with the GTRCAT for *CaM* (Bootstrap value: > 70%). *A. aurantiobrunneus* and *A. fructus* were used as outgroup. Numbers after taxa are isolate, specimen or strain numbers. The scale bar represents the number of nucleotide substitutions per site. The sequences obtained in this study are shown in bold. T = type sequence.

### Morphological observations

The colony morphologies of *A. latus* varied among strains (Fig. 2, Supplementary Fig. 1). On MEA, the growth was slower compared to that of *A. sublatus* and *A. spinulosporus* (Fig. 2B, Supplementary Fig. 1B, E, H, K, N, Q, T, W). In terms of colony appearance, *A. latus* was more similar to *A. spinulosporus* than to *A. sublatus*, as its colonies were covered with green conidia on OA and sometimes on CYA (Table 2 and Supplementary Fig. 1C, F, I, L, O, R, X). The microscopic of *A. latus* also closely resembled those of the other two species but ascospores of *A. spinulosporus* are echinulated (Table 2) [35, 36].

**Fig. 2.**
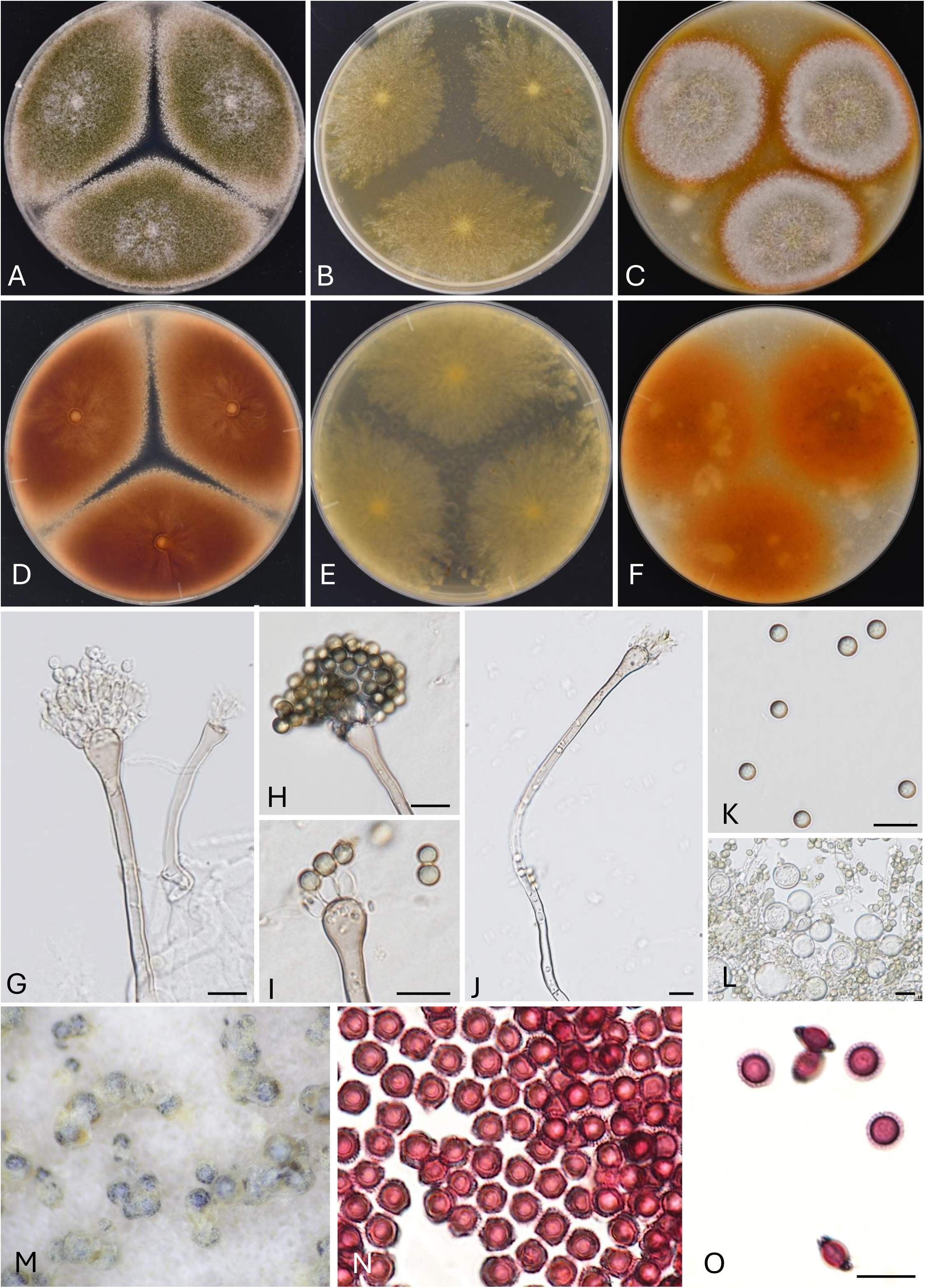
*A. latus* IFM 65329 colonies: obverse CYA 37 °C (A), MEA 37 °C (B), OA 25 °C (C), reverse: CYA 37 °C (D), MEA 37 °C (E), OA 25 °C, 7d (F), conidiophores (G–J), conidia (K), Hülle cells (L), ascomata (M), ascospores (N, O) Scale bars: G–L, N, O 10 μm.

**Table 2.**
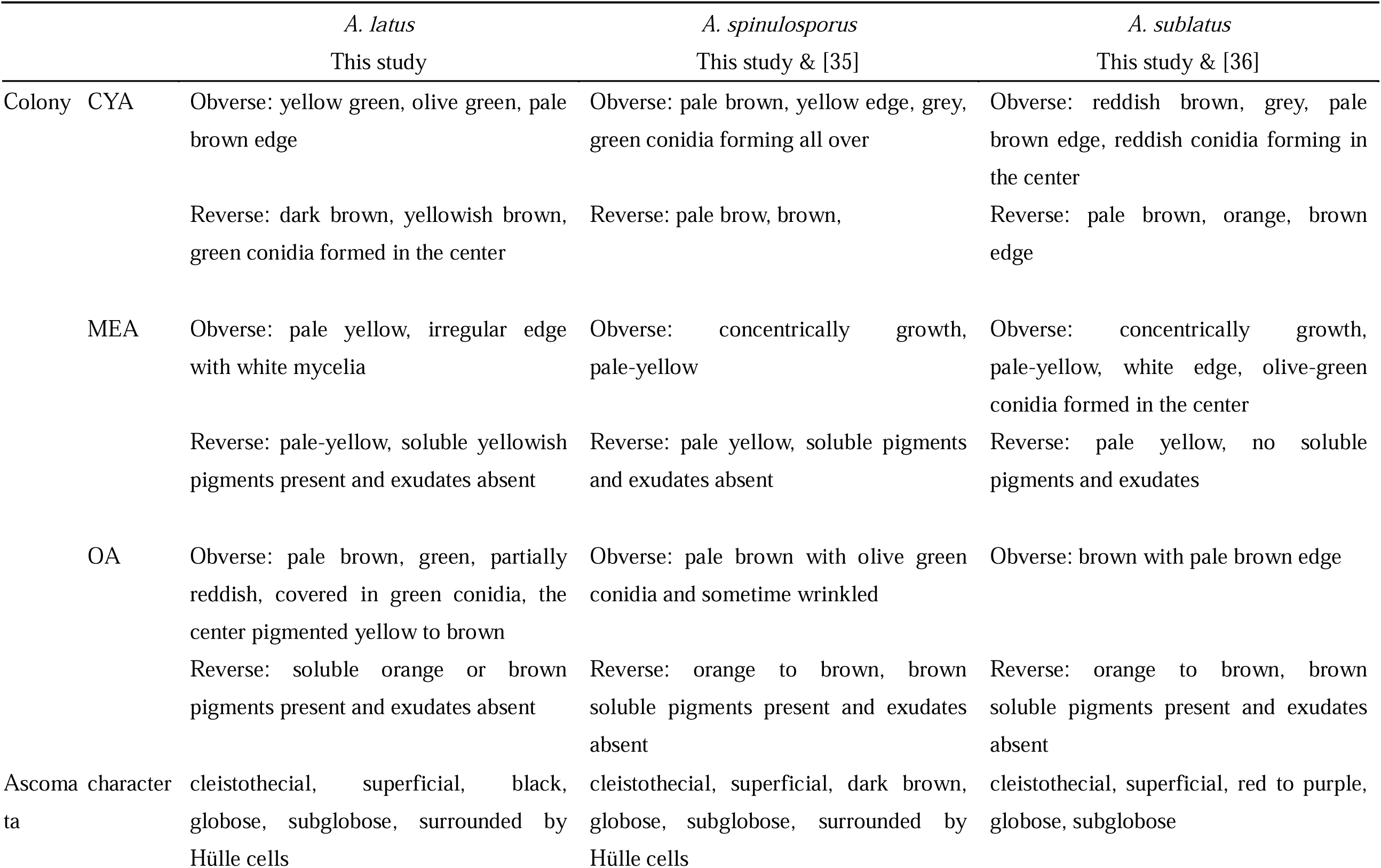
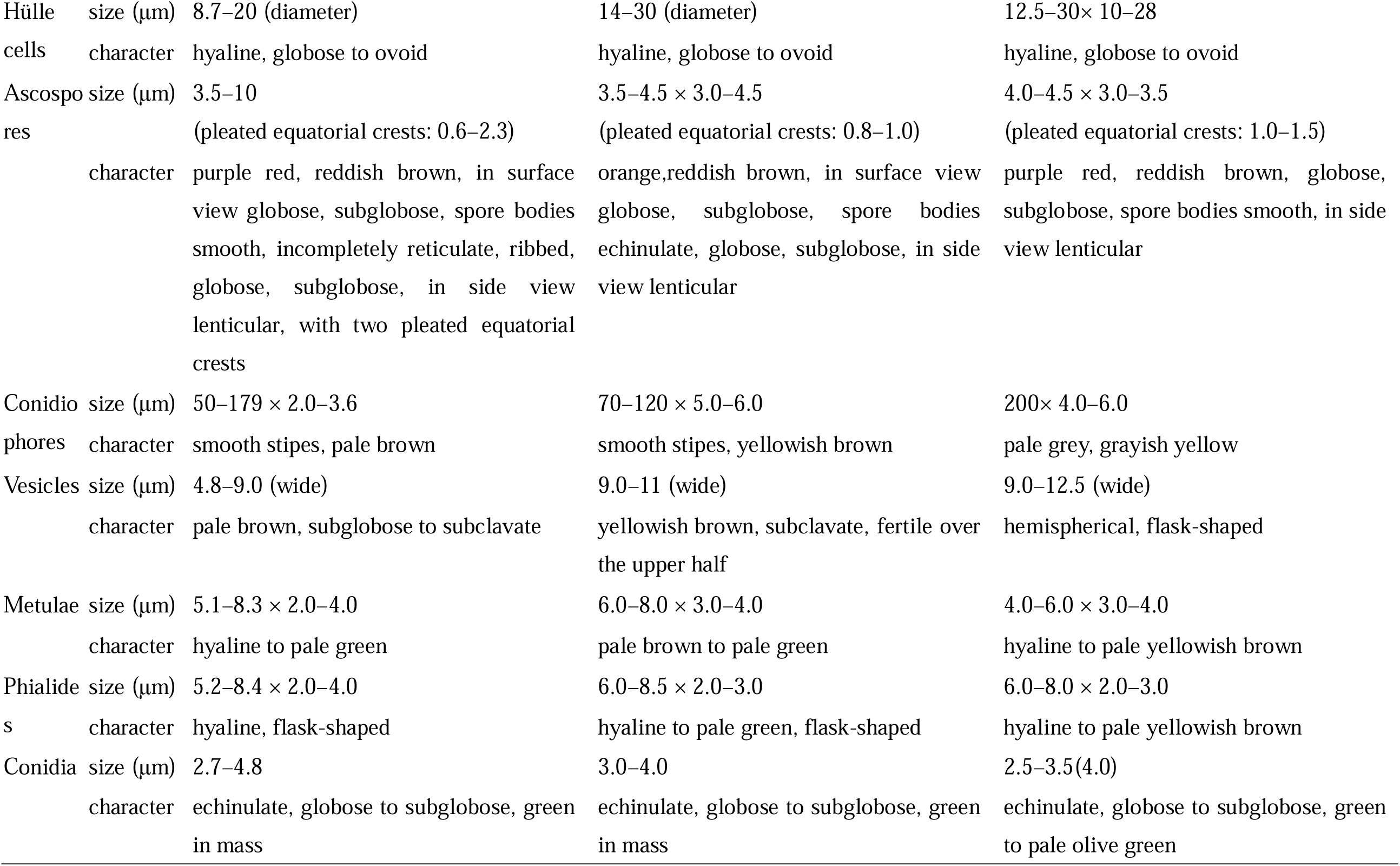
Morphological comparison of Aspergillus latus, A. spinulosporus and A. sublatus.

### Antifungal susceptibility testing

Antifungal susceptibility testing was conducted on seven *A. latus*, one *A. sublatus*, and three *A. spinulosporus* strains (Table 3). Reduced susceptibility to CPFG was observed in all tested strains, geometric mean MICs of 7.2 for *A. latus*, 16 for *A. sublatus*, and 5.0 for *A. spinulosporus*. Reduced susceptibility to AMPH-B was noted in *A. latus* and *A. sublatus*, while *A. spinulosporus* strains displayed low MICs, with geometric mean MICs of 4.4, 4.0, and 0.7, respectively. *Aspergillus latus* strains displayed low MICs to MCFG, whereas MICs of *A. sublatus*, and *A. spinulosporus* were elevated. Susceptibility to VRCZ and MCZ varied: reduced susceptibility to VRCZ was observed in *A. latus* IFM 66778 strain (MIC 2.0), and reduced susceptibility to MCZ was noted in *A. latus* strains IFM 61956 and IFM 65329 (MIC 2.0), as well as *A. sublatus*. All tested strains were susceptible to ITCZ.

**Table 3.** MIC values for antifungal drugs.

| Species name | Strain No. | MIC (µg/mL) |  |  |  |  |  |
| --- | --- | --- | --- | --- | --- | --- | --- |
|  |  | MCFG | CPFG | AMPH-B | ITCZ | VRCZ | MCZ |
| *Aspergillus latus* | IFM 61956 | 0.06 | 16 | 4 | 0.5 | 0.12 | 2 |
|  | IFM 63852 | 0.12 | 16 | 4 | 0.25 | 0.12 | 1 |
|  | IFM 64360 | 0.06 | 16 | 4 | 0.25 | 0.25 | 1 |
|  | IFM 65030 | 0.06 | 4 | 4 | 0.25 | 0.06 | 1 |
|  | IFM 65233 | 0.06 | 2 | 4 | 0.5 | 0.06 | 1 |
|  | IFM 65329 | 0.03 | 8 | 4 | 0.25 | 0.12 | 2 |
|  | IFM 66778 | 0.06 | 4 | 8 | 0.25 | 2 | 0.5 |
| *A. sublatus* | IFM 42029<sup>T</sup> | >16 | 16 | 4 | 0.25 | 0.12 | 2 |
| *A. spinulosporus* | IFM 66771 | 0.15> | 2 | 1 | 0.25 | 0.25 | 0.5 |
|  | IFM 61449 | >16 | 8 | 0.5 | 0.25 | 0.12 | 0.5 |
|  | IFM 64750 | >16 | 8 | 1 | 0.25 | 0.12 | 1 |

MCFG = micafungin, CPFG = caspofungin, AMPH-B = amphotericin B, ITCZ = itraconazole, VRCZ = voriconazole, MCZ = miconazole

## Discussion

Here, we report the allodiploid hybrid *A. latus* strains isolated in Japan. Although 30 strains have been found worldwide [8, 9], *A. latus* has not been reported in Japan as well as Asia. By screening 23 *A. spinulosporus* strains, we found seven clinical *A. latus* strains (30.4%). *A. latus* IFM 61956 was isolated in 2013, suggesting that this species might be more prevalent in the country. The pathogenicity of *A. latus* was shown to be comparable to that of *A. spinulosporus* and *A. nidulans* [8, 9]. In this study, the pathogenicity of *A. latus* was unclear, because *A. latus* strains could be colonized in most cases (Table 1). However, ongoing monitoring is necessary, as changes in pathogenicity or resistance profiles could impact treatment strategies in the future.

We conducted a phylogenetic analysis of *A. latus* (Fig. 1) with results indicating that it is derived from a hybridization between *A. spinulosporus* and a species related to *A. sublatus* and *A. quadrilineatus*. Morphological characters of seven *A. latus* strains, such as colony color and growth on MEA, were most stable among strains (Fig. 2B, Table 2, and Supplementary Fig. 1B, E, H, K, N, Q). The differences between *A. latus*, *A. spinulosporus* and *A. sublatus*, however, were slight, and given the variability observed in previous studies, species identification relying on these traits may be challenging (Supplementary Fig. 1) [9].

In terms of antifungal susceptibility, strains of *A. latus* found in Japan demonstrated increased susceptibility to ITCZ compared to overseas strains [8], while showing reduced susceptibility to CPFG, aligning with the previous research [8, 9] (Table 3). It has been reported that *A. spinulosporus* exhibits reduced susceptibility to CPFG compared to *A. nidulans* and other clinically relevant species in section *Nidulantes* [8, 37]. In this study, we confirmed that *A. spinulosporus* and *A. sublatus* showed reduced susceptibility to CPFG, suggesting that this characteristic in *A. latus* may be derived from these species. In addition, *A. latus* strains exhibited AMPH-B resistance. Although high AMPH-B MIC values have been reported in other *Aspergillus* species, especially *A. nidulans* and related species [8, 37, 38], the mechanisms underlying the resistance remain poorly understood. Increased catalase production has been implicated in AMPH-B resistance in *A. terreus* [39], suggesting that *A. latus* may produce high level of catalase.

In conclusion, our study provides significant insights into the ecology and clinical relevance of *A. latus*, a newly recognized hybrid pathogen and cryptic species of *A. nidulans*. We documented the first occurrence of this pathogen in Japan, demonstrating its prevalence and potential role as a causative agent of aspergillosis since 2013. The comprehensive phenotyping, including morphological and genetic analysis, underscored the diagnostic challenges due to its similarities with *A. spinulosporus* and *A. sublatus*. Moreover, antifungal susceptibility testing revealed reduced susceptibility to certain antifungal agents, highlighting the need for accurate identification to inform effective treatment strategies. Our findings add to the growing body of knowledge about the distribution and pathogenic characteristics of *A. latus* and emphasize the importance of continued surveillance and accurate diagnostics in clinical mycology.

## Acknowledgements

We thank Dr Takayuki Arazoe (Tokyo University of Science) for providing us with the pMK-dGFP plasmid. We thank Junko Ito and Emi Shirai for technical assistance with the experiments. We are also grateful to Machiko Zen for assistance with the data analysis.

## Statements & Declarations

### Author Contributions

Saho Shibata, Vit Hubka, and Hiroki Takahashi contributed to the study conception and design. Material preparation, data collection and analysis were performed by Saho Shibata, Momotaka Uchida, Sayaka Ban, Kamei Katsuhiko, Akira Watanabe, Vit Hubka, and Hiroki Takahashi. The draft of the manuscript was written by Saho Shibata, Momotaka Uchida, Takashi Yaguchi, Vit Hubka and Hiroki Takahashi, and all authors commented on the manuscript. All authors read and approved the final manuscript.

### Ethical approval

No ethical approval is required

