## Supplementary Figure 1 for "*Aspergillus latus*: a cryptic causative agent of aspergillosis emerging in Japan"

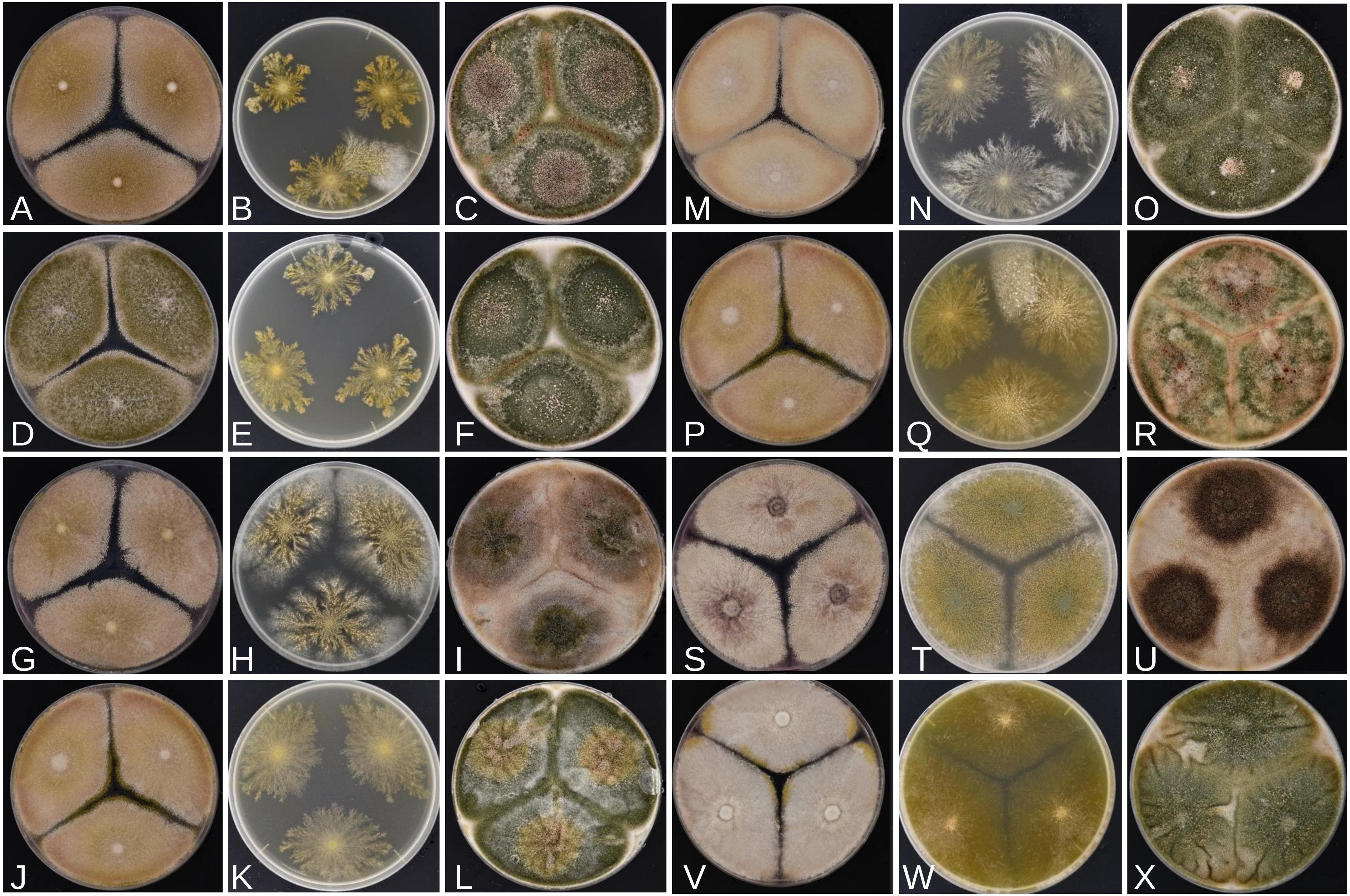


**Supplementary Fig. 1**. *Aspergillus latus* IFM 61956 (A–C), IFM 63852 (D–F), IFM 64360 (G–I), IFM 65030 (J–L), IFM 65233 (M–O), IFM 66778 (P–R). *A. sublatus* IFM 42029$^{T}$ (S–U). *A. spinulosporus* IFM 66771 (V–X). Colonies : 37 ° C, 7d, on CYA (A, D, G, J, M, P, S, V), MEA (B, E, H, K, N, Q, T, W), OA (C, F, I, L, O, R, U, X).
